# Environmental DNA enables rapid detection of invasive coypu and complements camera trapping

**DOI:** 10.64898/2026.09.11.749317

**Authors:** Cyrielle Ballester, Simon Lacombe, Sébastien Devillard, Louise D’Hollande, Yann Raulet, Vincent Sablain, Geoffrey Didier, Raphaël Mathevet, Claude Miaud, Clément Oyon, Sébastien Richarte, Serge Rouvière, Alice Valentini, Nathalie Vazzoler-Antoine, Olivier Gimenez

**Affiliations:** CEFE, CNRS, EPHE, IRD, Université de Montpellier, Montpellier, France; Universite Claude Bernard Lyon 1, LBBE, UMR 5558, CNRS, VAS, Villeurbanne, 69100, France; EPTB Symbo, Boulevard de la Démocratie, Mauguio, France; Service Nature Observatoire et Territoire, Direction Paysage et Biodiversité, Ville de Montpellier, Montpellier, France; EPTB Lez (Syndicat du Bassin du Lez-SYBLE), Prades-le-Lez, France; Association Fiber Nature, Montpellier, France; EPTB Vidourle, Sommières, France; Métropole de Montpellier, Service GEMAPI, Montpellier, France; SPYGEN, Le Bourget-du-Lac cedex, France

**Author notes:** Co-first authors (Cyrielle Ballester, Simon Lacombe).

**Keywords:** biological invasions, camera trapping, detection probability, environmental DNA metabarcoding, invasive species monitoring, Myocastor coypus, One Health

## Abstract

Effective management of invasive species requires surveillance methods that reliably detect populations while minimizing field effort. Environmental DNA (eDNA) offers a potentially rapid alternative to conventional monitoring, but direct comparisons with other non-invasive methods remain limited for invasive mammals.

Here, we compared eDNA water sampling and camera trapping for detecting established populations of coypu (*Myocastor coypus*), a widespread invasive semi-aquatic rodent. Using replicated data collected across Mediterranean river systems in southern France, we estimated detection probabilities conditional on coypu presence using Bayesian generalized linear mixed models and quantified the sampling effort required to achieve high cumulative detection probability.

Detection probability was 93.0% (95% credible interval: 72.1-98.9%) for a single eDNA water sample, compared with 52.8% (22.6-91.1%) for 30 camera-trap days, with a 96.9% posterior probability that eDNA detection was higher. Two eDNA samples were sufficient to achieve a median cumulative detection probability above 95%, compared with four 30-day camera-trapping periods. Detection also varied at different spatial scales, with eDNA detectability varying more among catchments and camera-trap detectability being heterogeneous among local camera locations.

These results show that eDNA provides a rapid and reliable approach for confirming the presence of established coypu populations, and likely to detect the species in newly recolonized areas, while camera traps provide complementary information on activity, behaviour and habitat use. Combining rapid eDNA screening with targeted camera trapping may therefore provide an efficient surveillance strategy for invasive semi-aquatic mammals.

## Introduction

Invasive alien species are major drivers of ecological change and generate substantial economic, social and public-health costs (Pyšek et al. 2020, Roy et al. 2024). Their effective management depends critically on the ability to determine where populations occur, track changes in their distribution, and evaluate the outcomes of control actions. Yet surveillance of invasive species remains challenging because animals are imperfectly detected, and monitoring methods must often balance reliability against the time, effort and resources required for field sampling. These challenges are particularly important for invasive rodents, whose impacts can affect ecosystems, infrastructure, agriculture and human health (Diagne et al. 2023).

The coypu (*Myocastor coypus*) provides a particularly relevant example. Native to South America, the species has established widespread invasive populations across Europe, including France, following introductions associated with fur farming (Schertler et al. 2020, Bonnet et al. 2023). Coypus can cause ecological and economic damage through grazing and burrowing, and their management involves control operations in many parts of their introduced range. Beyond these ecological and management concerns, coypus are also associated with important sanitary risks. In particular, they can act as healthy carriers of pathogenic *Leptospira*, the bacteria responsible for leptospirosis in humans (Michel et al. 2001), placing their surveillance at the intersection of invasive species management and One Health.

Several approaches can be used to monitor coypu populations, ranging from direct observations and trapping to non-invasive methods such as camera trapping and environmental DNA (eDNA). These approaches provide different types of information and involve different logistical constraints. Trapping can provide information on individuals and can be integrated into control operations, but requires substantial field effort and, when used for inference on population abundance, appropriate methods to account for imperfect detection (Gimenez 2025). Camera traps offer a non-invasive alternative and can provide rich information on species occurrence, activity and behaviour (Pollet et al. 2025, Bruce et al. 2025). For example, year-round camera trapping of an urban coypu population revealed marked temporal variation in activity and a diverse behavioural repertoire, information that could directly inform the timing and targeting of control operations (Viviano et al. 2025). However, cameras typically require equipment to remain deployed for weeks or months and subsequent processing of images or videos. By contrast, eDNA sampling can provide evidence of species presence from water samples collected during a single site visit, potentially allowing large numbers of sites to be surveyed over a much shorter period.

The potential of eDNA for coypu surveillance has already been demonstrated. A species-specific qPCR assay successfully detected coypu from water samples at sites of known presence in the United States, with detection strongly influenced by sampling methodology (Mangan et al. 2023). In particular, filtration of relatively large volumes of water produced substantially higher detection than direct sampling of small water volumes. Subsequent field validation showed that eDNA could also provide a useful tool for detecting reinvasion and monitoring populations following eradication, when conventional surveillance may become particularly resource-intensive (Coster 2024). Together, these studies establish eDNA as a promising surveillance tool for coypu, but its detection performance relative to another widely applicable non-invasive method such as camera trapping remains poorly quantified under a common sampling design (see Croose et al. 2023 for the European mink *Mustela lutreola*, Girolamo et al. 2024 for the American mink *Neogale vison*, Lacombe et al. 2026 for the Eurasian otter *Lutra lutra*).

Here, we compared eDNA sampling and camera trapping for detecting coypu populations at the same sampling sites. Specifically, we quantified and compared detection probabilities, and evaluated the sampling effort required by each approach to achieve high detection probability. Our study originates from a comparison of non-invasive methods for monitoring the European otter, in which water eDNA metabarcoding revealed frequent and abundant detections of coypu DNA (Lacombe et al. 2026). This provides an opportunity to evaluate the performance of eDNA metabarcoding for an abundant invasive mammal under a common spatial sampling design with camera trapping. Rather than asking which method is universally superior, our objective was to assess their respective strengths for surveillance of established coypu populations and their potential complementarity for the management of invasive species.

## Methods

### Study area

The study was conducted in the Gard and Hérault departments of southern France and encompassed four Mediterranean river systems: the Lez River, the Mosson River, the Étang de l’Or catchment, and the Vidourle River. All four river systems experience a Mediterranean climate characterized by hot, dry summers and episodes of intense rainfall, particularly in autumn, resulting in strong seasonal and short-term variation in river discharge. Riparian vegetation is broadly characteristic of Mediterranean lowland rivers, and all four systems are modified to varying degrees by artificial structures and river engineering (Lacombe et al. 2026).

The Lez and Mosson rivers flow through the Montpellier metropolitan area and are therefore strongly influenced by urbanization, although both retain sections of more natural riparian habitat. The Lez, a 29.6-km river, is particularly modified by numerous weirs, bridges and other artificial structures. The Étang de l’Or catchment consists of a coastal lagoon connected to the Mediterranean Sea, together with several tributaries, artificial canals and extensive wetlands. Its landscape ranges from Mediterranean scrubland upstream to a mosaic of agricultural, urban and wetland habitats closer to the lagoon. The Vidourle River extends from the Cévennes foothills to the Mediterranean Sea, flowing through relatively natural and forested landscapes upstream and predominantly rural and agricultural areas further downstream. All four river systems support established coypu populations.

### Sampling design

Data were originally collected as part of a multi-method survey designed to compare approaches for detecting Eurasian otters (Lacombe et al. 2026). Because the eDNA metabarcoding approach simultaneously detected mammalian species other than the focal species, the resulting data also provided replicated detections of coypu. Here, we use these data to compare eDNA and cameratrap detection of coypu across all four catchments.

Fieldwork was conducted over approximately two months in each catchment, in late autumn for the Lez (2023) and Vidourle (2024), and in early spring for the Étang de l’Or (2024) and Mosson (2025). Within each catchment, camera trapping and eDNA sampling therefore covered the same study period, although sampling periods differed among catchments. We surveyed 5-8 sites per catchment, for a total of 28 sites. Each site consisted of a 600-m river stretch, with neighbouring sites separated by at least 1 km to reduce spatial dependence. Sites were selected primarily according to river accessibility and the feasibility of implementing the different monitoring protocols.

At each site, coypu occurrence was assessed using two independent non-invasive approaches: continuous camera trapping and three replicate eDNA water samples collected during a single sampling occasion. Further details of the overall sampling design are provided in Lacombe et al. (2026).

### Camera trapping

Two camera traps (Browning SPEC OPS ELITE HP5) were deployed at each site, generally near the upstream and downstream ends of the 600-m river stretch. Cameras were positioned close to the river at locations expected to maximize encounters with otters, including riverbanks, animal trails, gravel bars and artificial structures such as bridges or weirs. They operated continuously using motion sensors and recorded 30-s videos following each trigger.

Cameras remained in the field for approximately two months and were checked periodically to replace batteries and memory cards. All videos were manually reviewed and species were identified following the protocol described in Lacombe et al. (2026). Coypu records were subsequently aggregated into 24-h sampling occasions, yielding daily detection/non-detection histories for each camera.

### Environmental DNA sampling and analysis

Environmental DNA sampling and laboratory analyses followed the protocol described by Lacombe et al. (2026). Briefly, water was sampled once at each site near the downstream end of the 600-m river stretch. Three independent filtration replicates were collected at the same location and on the same day. For each replicate, approximately 30L of river water were filtered over 30min using a peristaltic pump (Vampire sampler, Bürkle, Germany) and sterile disposable tubing connected to a VigiDNA 0.45-µm cross-flow filtration capsule (SPYGEN, Le Bourget-du-Lac, France). Following filtration, capsules were drained, filled with 80 mL of CL1 preservation buffer (SPYGEN), and stored at room temperature until DNA extraction following the protocol described in Pont et al. (2018).

Following DNA extraction, samples were screened for PCR inhibition (Biggs et al. 2015) and diluted when necessary. Metabarcoding targeted fragments of the mitochondrial 12S region. We used the Mamm01 primers (Taberlet et al. 2018) for samples from the Lez and Bassin de l’Or catchments, and the vertebrate V05 primers (Riaz et al. 2011) for samples from the Vidourle and Mosson catchments. Twelve PCR replicates were performed for each filtration capsule, with sample-specific tagged primers, and amplification products were pooled before library preparation to achieve a theoretical sequencing depth of 300 000 reads per samples and sequenced on an Illumina NextSeq platform. Extraction and amplification negative controls were included throughout the procedure and sequenced in parallel.

Sequence processing and taxonomic assignment followed Lacombe et al. (2026) using OBITools tools (Boyer et al. 2016), with taxonomic assignments made against GenBank release 247. Low-frequency and low-quality sequences were removed, and additional filtering was applied to account for tag-jumping and index-hopping. Further details on DNA extraction, amplification, sequencing, bioinformatic filtering and quality-control procedures are provided in Lacombe et al. (2026). .The species found in the negative controls were human and domestic animals (e.g. chicken, turkey, pig, dog and/or cat). After the filtering pipeline, the extraction and PCR negative controls were completely clean, and no sequence reads remained in those samples.

### Statistical analyses

We estimated detection probabilities for each method conditional on coypu presence, assuming that coypu were present at all surveyed sites. Detection/non-detection data were analyzed using generalized linear mixed models (GLMMs) with a Bernoulli distribution and a logit link. We fitted separate intercept-only models for eDNA and camera-trap data. Both models included a catchment-level random intercept to account for variation in detectability among river systems. The eDNA model additionally included a site-level random intercept to account for local variation among sampling sites, whereas the camera-trap model included a camera-level random intercept to account for heterogeneity among individual camera locations.

For eDNA, the sampling replicate was a single water sample, and detection probability was therefore estimated per water sample. For camera trapping, the sampling replicate was initially defined as a camera-day to accommodate variation in camera deployment duration. Daily detection probability, p_CT_, was subsequently converted to the probability of detecting coypu over 30 camera-days as 1 − (1 − p_CT_)^30^. This provided a more meaningful unit of camera-trapping effort for comparison with eDNA sampling (Lacombe et al. 2026).

For each method, we used posterior distributions to estimate the average detection probability and catchment-specific detection probabilities. We also quantified the relative contribution of catchment- and local-level heterogeneity to variation in detectability using variance partition coefficients. Finally, we derived cumulative detection curves describing the probability of detecting coypu at least once after k sampling replicates, p_k_ = 1 − (1 − p)^k^, where a replicate corresponded to one water sample for eDNA and 30 camera-days for camera trapping. From these curves, we calculated the minimum number of replicates required to achieve a cumulative detection probability of at least 95%.

Models were fitted in a Bayesian framework using R v4.6.1 (R Core Team 2025) and the *brms* package (Bürkner 2017). We assigned a weakly informative N(0,2.5) prior to the intercept and Exponential(1) priors to the standard deviations of random effects. Models were run for 5,000 iterations, including 1,000 warm-up iterations, across two Markov chains. Convergence was assessed by visual inspection of trace plots and by ensuring that R-hat ≤1.01 for all parameters.

### Results

Coypu detection probability was higher using eDNA than camera trapping. Coypu were detected by camera traps at 22 of the 28 surveyed sites (3/5 in the Mosson, 5/7 in the Lez, 6/8 in the Or catchment and 8/8 in the Vidourle), compared with 27 of 28 sites using eDNA (5/5, 7/7, 7/8 and 8/8, respectively). The estimated probability of detecting coypu from a single eDNA water sample was 93.0% (95% CrI: 72.1-98.9%), compared with 52.8% (95% CrI: 22.6-91.1%) over 30 cameratrap days. The posterior probability that detection was higher with eDNA than with 30 days of camera trapping was 96.9%.

This difference was consistent across catchments (Fig. 1, left panel). Catchment-specific eDNA detection probabilities ranged from 82.4% (95% CrI: 58.8-95.5%) in the Or catchment to 97.3% (87.8-99.9%) in the Vidourle. In contrast, 30-day camera-trap detection probabilities ranged from 40.3% (7.0-77.9%) in the Mosson to 60.2% (26.6-94.0%) in the Vidourle.

**Figure 1.**
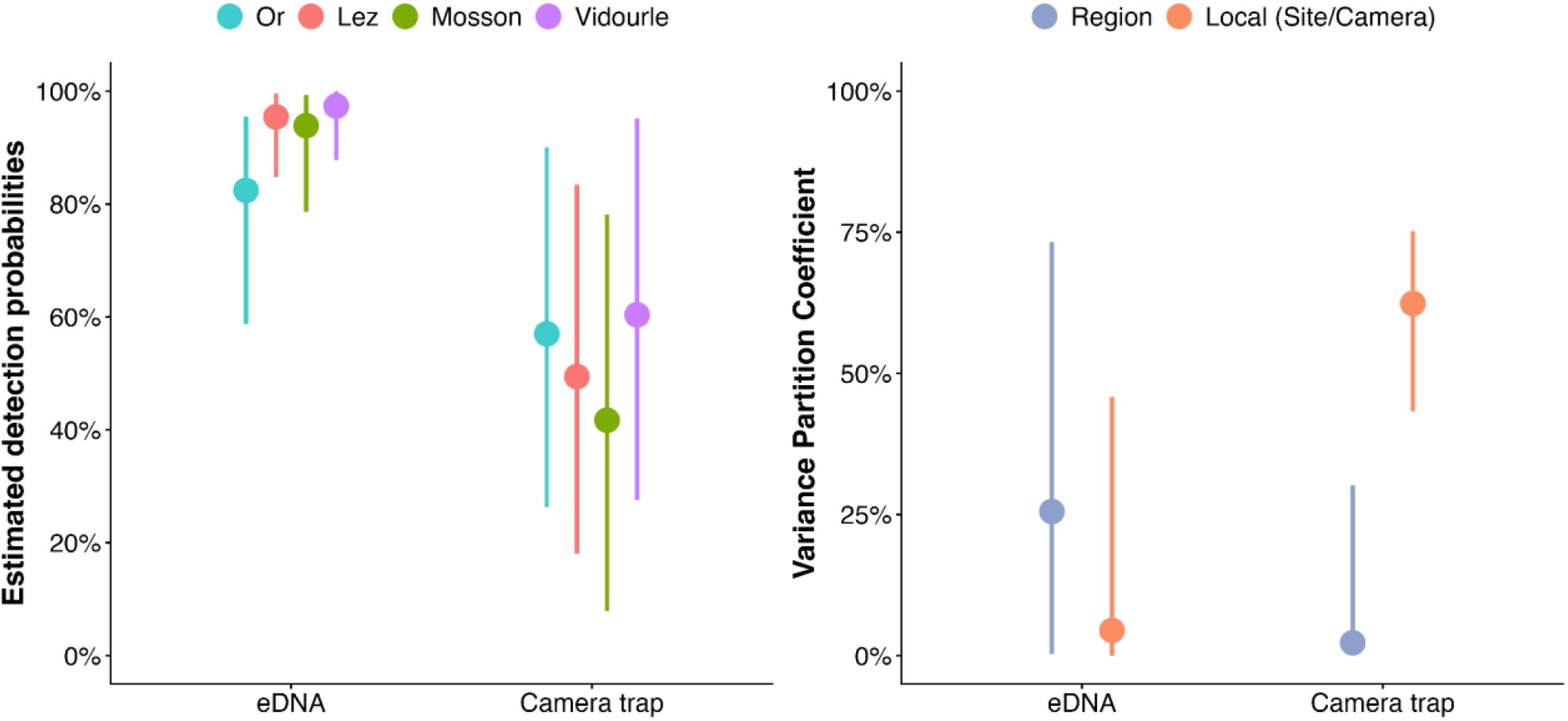
Spatial variation in coypu detection probability using environmental DNA (eDNA) and camera trapping. Left: Estimated detection probabilities for the four study catchments (Or, Lez, Mosson and Vidourle). For eDNA, probabilities correspond to a single water sample; for camera trapping, probabilities correspond to 30 camera-trap days. Right: Variance partition coefficients describing the proportion of total variance attributable to catchment-level and local-level (site for eDNA; camera for camera trapping) heterogeneity. In both panels, points show posterior medians and error bars indicate 95% credible intervals.

The spatial structure of variation in detectability also differed markedly between monitoring methods (Fig. 1, right panel). For camera trapping, local heterogeneity among cameras accounted for a median 61.8% (95% CrI: 42.2-75.1%) of the total variance, whereas differences among catchments accounted for only 2.5% (0.004-31.5%). For eDNA, local variation among sampling sites was lower, accounting for 4.5% (0.01-45.6%) of total variance, while catchment-level variation accounted for 24.7% (0.2-74.0%).

The higher per-replicate detection probability of eDNA translated into a rapid increase in cumulative detection probability with sampling effort (Fig. 2). Two eDNA water samples were sufficient to exceed a median cumulative detection probability of 95%, with the estimated number of samples required to reach this threshold ranging from one to three across the 95% credible interval. In comparison, camera trapping required a median of four 30-day sampling periods to exceed 95% cumulative detection probability, with greater uncertainty (95% CrI: 2-12 months).

**Figure 2.**
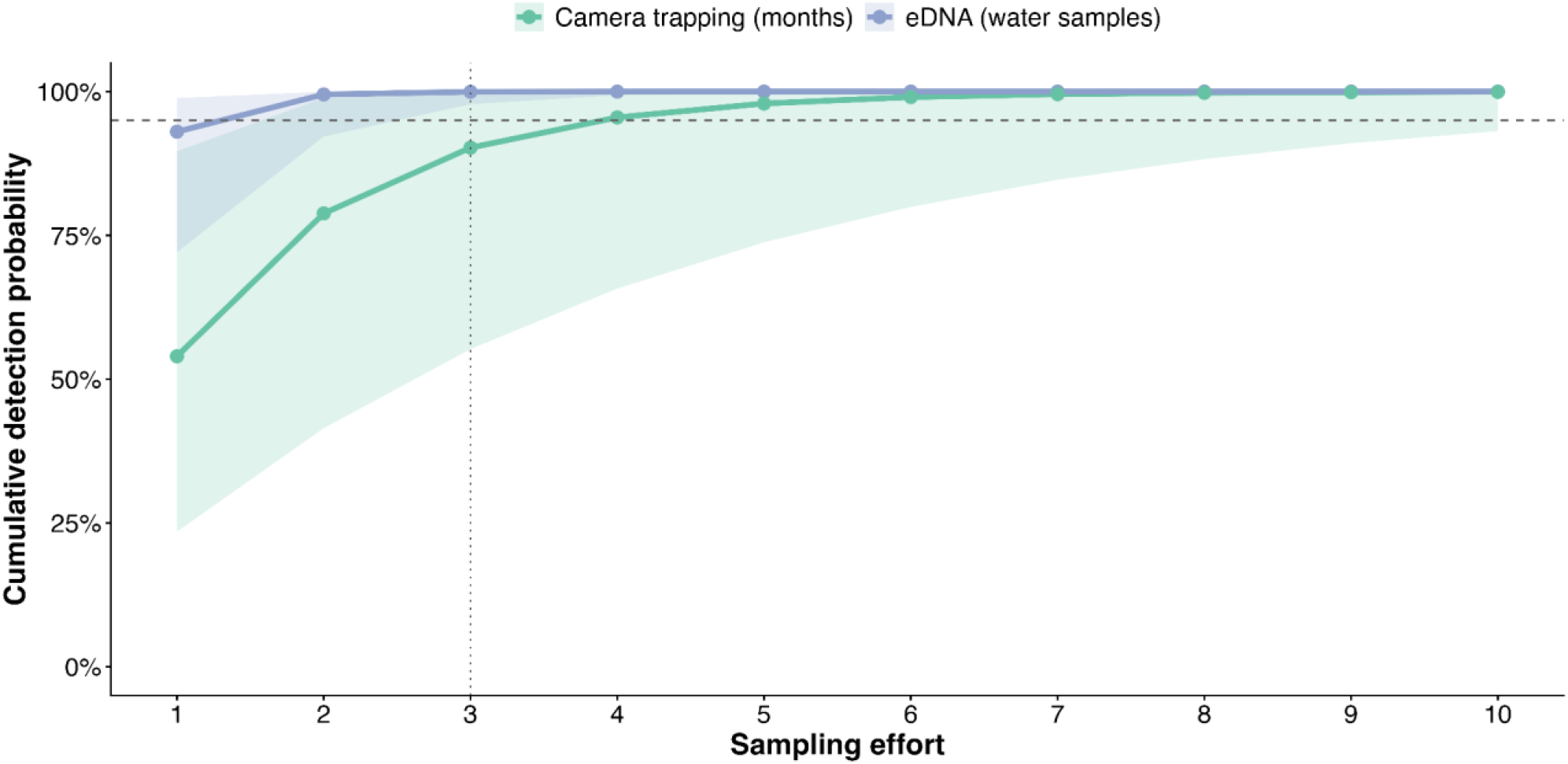
Cumulative probability of detecting coypu as a function of sampling effort using environmental DNA (eDNA) and camera trapping. Sampling effort corresponds to the number of water samples for eDNA and the number of 30-day deployment periods for camera trapping. Lines show posterior medians and shaded areas indicate 95% credible intervals. The horizontal dashed line indicates a cumulative detection probability of 95%, and the vertical dotted line indicates three sampling replicates, corresponding to the maximum number of eDNA water samples collected per site.

## Discussion

Our results highlight the high capacity of environmental DNA for detecting established coypu populations. A single eDNA water sample had a high probability of detecting coypu, and only two samples were required to reach a median cumulative detection probability above 95%. Camera trapping required substantially greater sampling effort to achieve comparable detection. These findings reinforce previous evidence that eDNA can provide an effective surveillance tool for coypu (Mangan et al. 2023, Coster 2024), while our direct comparison shows that this advantage can be substantial relative to camera trapping when the primary objective is simply to establish species presence. However, rather than identifying a universally superior monitoring method, we emphasize that method choice should depend on the ecological or management question being addressed.

### eDNA provides rapid and reliable detection of coypu

The high detection probability obtained from individual water samples suggests that eDNA is particularly well suited to rapid surveillance of established coypu populations. This is consistent with Mangan et al. (2023), who detected coypu at all known-presence sites using a single filtration sample with qPCR and showed that sampling methodology can strongly influence detection. Coster (2024) similarly emphasized the operational advantage of eDNA for surveying multiple sites efficiently and its potential contribution to post-eradication monitoring. This advantage is especially relevant in terms of field deployment when sampling many sites: eDNA samples can be collected during a single visit, whereas cameras must remain deployed for extended periods and subsequently be retrieved. Both approaches nevertheless require substantial post-field processing, involving laboratory and bioinformatic analyses for eDNA and image or video processing for camera trapping.

Interestingly, coypu DNA was initially encountered as a highly abundant signal in an eDNA survey primarily designed to detect Eurasian otters (Lacombe et al. 2026). What represented background DNA when targeting a relatively rare carnivore became a strong surveillance signal for an established invasive rodent. This also illustrates an advantage of metabarcoding: samples collected for one monitoring objective can simultaneously provide information on other species of ecological, management or public health interest. This broader taxonomic coverage may, depending on the marker and protocol used, come at some cost in sensitivity compared with species-specific approaches such as qPCR (McColl-Gausden et al. 2023). However, this trade-off is not universal, as metabarcoding protocols can achieve detection rates comparable to those of species-specific approaches in some systems (Condachou et al. 2024).

The spatial structure of detectability also differed between methods. Camera-trap detection varied predominantly among individual camera locations, consistent with a strong influence of camera placement and local animal movements (Burton et al. 2015, Sanders et al. 2024, Croose et al. 2025). Importantly, camera locations in our study were selected primarily to maximize encounters with otters, the focal species of the original survey (Lacombe et al. 2026), rather than coypu. Although both species are semi-aquatic and use similar riparian habitats, species-specific differences in behaviour and fine-scale habitat use may influence the locations at which encounters with cameras are most likely. In particular, camera-trap detection of coypu may have been lower than under a survey specifically designed for this species. In contrast, eDNA detection varied little among sites within catchments but more strongly among catchments, suggesting that water sampling may integrate species presence over a broader spatial scale and be less sensitive to the precise sampling location (Deiner et al. 2016, Harper et al. 2019, Lyet et al. 2021, Altermatt et al. 2025). The greater variation among catchments could reflect differences in coypu abundance, but also differences in environmental and hydrological conditions affecting eDNA release, transport, dilution or persistence. Our study was not designed to distinguish among these potential sources of catchment-level variation.

### Different tools answer different ecological questions

Despite its greater detection sensitivity, eDNA should not be viewed as a replacement for camera trapping. eDNA is particularly efficient for answering whether a species is present, whereas cameras provide temporally explicit observations that can inform activity patterns, behaviour and habitat use. Viviano et al. (2025), for example, used camera trapping to characterize coypu activity rhythms and behavioural repertoires and showed how this information could help target control operations in space and time.

The two methods therefore provide complementary information (Croose et al. 2023, Girolamo et al. 2024). This complementarity suggests a simple surveillance strategy: eDNA could be used for rapid screening across many sites, followed by targeted camera trapping at positive or priority sites where richer ecological information is required. The spatial interpretation of detections also differs between methods. Whereas a camera-trap detection provides direct evidence that an animal occurred at the camera location, eDNA transported by flowing water may originate some distance upstream from the sampling point (Wood et al. 2021). Consequently, using eDNA to delineate coypu distribution at fine spatial scales requires a sampling design that accounts for the structure and hydrology of the river network.

### Implications for invasive species management and One Health

Rapid and reliable detection is central to the management of invasive species, from mapping occupied areas to targeting control and evaluating management interventions (Yackel Adams et al. 2024, Gimenez 2025). This is especially important for coypu because small populations can rapidly increase and populations have re-established following intensive control programmes (Bonnet et al. 2023). Coster (2024) specifically proposed eDNA as a promising approach for post-eradication surveillance, when resources for intensive field monitoring may decline but failure to detect reinvasion can compromise previous management investment. Our results suggest that eDNA could provide an efficient tool for these purposes, although its performance at low population densities, particularly following intensive control, remains to be evaluated (Dejean et al. 2012, Davis et al. 2023).

More broadly, our results suggest how eDNA and camera trapping could be combined within an operational surveillance strategy for coypu. In areas with established populations, eDNA could provide a rapid first-line screening tool to map broad patterns of occurrence across river networks, followed by targeted camera trapping where information on activity, behaviour or fine-scale habitat use is needed to guide management. At invasion fronts or following intensive control, where populations are expected to occur at lower densities, combining methods may help reduce the risk of false negatives, although the relative performance of eDNA and camera trapping under these conditions remains to be evaluated.

Coypu surveillance also has a One Health dimension because the species can carry pathogenic *Leptospira*. Reliable information on coypu distribution may therefore contribute to assessing potential zoonotic risk in freshwater systems. Metabarcoding is particularly attractive in this context because the same samples can simultaneously provide information on multiple species (including pathogens), although detecting coypu alone does not provide information on infection or transmission risk.

### Limitations and conclusion

Our results should not be generalized to all invasive mammals, or even all stages of coypu invasion. Coypu were established in our study systems and are strongly associated with aquatic habitats, providing favourable conditions for releasing DNA directly into the sampled environment. The effectiveness of water-based eDNA surveillance may therefore depend strongly on species ecology and, in particular, on how frequently animals interact with the aquatic environment. Comparisons with other semi-aquatic mammals support this idea: for American mink, for example, eDNA and camera trapping showed broadly similar detection performance (Girolamo et al. 2024), while camera trapping outperformed eDNA for Eurasian otters under the same sampling framework as our study (Lacombe et al. 2026). Thus, the particularly high eDNA detectability observed here may partly reflect the close association of coypu with the aquatic environment. Detectability by both eDNA and camera trapping is likely to decline at lower population densities, potentially increasing the sampling effort required for reliable detection. For eDNA specifically, the number of samples required can increase when DNA concentrations are low (Mangan et al. 2023). Evaluating the relative performance of the two methods at low population densities will therefore be particularly important for early detection at invasion fronts and for post-control or post-eradication surveillance (Davis et al. 2023), arguably the situations in which management decisions are most sensitive to false negatives.

The contrast with results obtained for Eurasian otters under the same sampling framework is particularly informative: camera trapping provided otter detections at more sites than eDNA, whereas the opposite pattern emerged here for coypu. This suggests that monitoring performance depends not only on the method itself, but also on species abundance, ecology and behaviour. The higher eDNA detectability observed for coypu could therefore reflect several, potentially interacting factors, including differences in population density, biomass, use of the aquatic environment and rates of DNA release, which our study was not designed to disentangle.

Rather than supporting a single best monitoring method, our results therefore argue for surveillance strategies tailored to the target species and management objective. For established coypu populations, eDNA appears particularly well suited to rapid screening, with camera trapping providing a complementary tool when finer-scale ecological information is needed.

## Acknowledgements

We thank all the people who supported us in the field, in particular Lucas Buffan, Matthieu Rigal, Axelle Scamps, Tatiana Tronel, Louis Barbu and the members of the HAIR team at CEFE.

## Data, scripts, code, and supplementary information availability

Data and codes are available online https://doi.org/10.5281/zenodo.21878553

